# Patterns of individual variation in visual pathway structure and function in the sighted and blind

**DOI:** 10.1101/065441

**Authors:** Geoffrey K. Aguirre, Ritobrato Datta, Noah C. Benson, Sashank Prasad, Samuel G. Jacobson, Artur V. Cideciyan, Holly Bridge, Kate E. Watkins, Omar H. Butt, Alexsandra S. Dain, Lauren Brandes, Efstathios D. Gennatas

## Abstract

Many structural and functional brain alterations accompany blindness, with substantial individual variation in these effects. In normally sighted people, there is correlated individual variation in some visual pathway structures. Here we examined if the changes in brain anatomy produced by blindness alter the patterns of anatomical variation found in the sighted. We derived eight measures of central visual pathway anatomy from a structural image of the brain from 59 sighted and 53 blind people. These measures showed highly significant differences in mean size between the sighted and blind cohorts. When we examined the measurements across individuals within each group we found three clusters of correlated variation, with V1 surface area and pericalcarine volume linked, and independent of the thickness of V1 cortex. These two clusters were in turn relatively independent of the volumes of the optic chiasm and lateral geniculate nucleus. This same pattern of variation in visual pathway anatomy was found in the sighted and the blind. Anatomical changes within these clusters were graded by the timing of onset of blindness, with those subjects with a post-natal onset of blindness having alterations in brain anatomy that were intermediate to those seen in the sighted and congenitally blind. Many of the blind and sighted subjects also contributed functional MRI measures of cross-modal responses within visual cortex, and a diffusion tensor imaging measure of fractional anisotropy within the optic radiations and the splenium of the corpus callosum. We again found group differences between the blind and sighted in these measures. The previously identified clusters of anatomical variation were also found to be differentially related to these additional measures: across subjects, V1 cortical thickness was related to cross-modal activation, and the volume of the optic chiasm and lateral geniculate was related to fractional anisotropy in the visual pathway. Our findings show that several of the structural and functional effects of blindness may be reduced to a smaller set of dimensions. It also seems that the changes in the brain that accompany blindness are on a continuum with normal variation found in the sighted.

## Introduction

Blindness produces a well-documented set of structural changes along the visual pathway of the human brain. The optic nerves, lateral geniculate nucleus, and pericalcarine white matter are reduced in volume [1–4], as is the splenium of the corpus callosum, which connects the hemispheres of the visual cortex [5, 6]. While the surface area of primary visual cortex is reduced [7, 8], the gray matter layer of this area is thickened [7–10], perhaps reflecting altered synaptic pruning during cortical maturation [11, 12].

These mean differences between blind and sighted groups are present against a backdrop of substantial individual variation in visual pathway anatomy. In the sighted, there is a twoto-three fold variation in the size of visual pathway structures, and the relative size of some of these structures has been found to co-vary across individuals [13]. This diversity (in the case of occipital surface area) is influenced by both genetic and environmental factors [14–16] and is related to differences in visual acuity [17, 18]. Some elements of the visual pathway co-vary across individuals, while other elements vary independently [19, 20], suggesting separate factors influence these patterns of anatomical variation. In the blind, there is marked heterogeneity in the magnitude of brain alterations induced by vision loss. This variation has been linked to the age of blindness onset [5, 10, 21], although other factors—such as the degree or form of ophthalmologic injury—could contribute as well.

Here we characterize the pattern of shared and independent anatomical variation that is present along the visual pathway. Our first goal was to identify how individual variation in the size of anatomical features cluster together across individuals. Next, we examined if this pattern of clustering is the same across sighted and blind people. Similar patterns of variation in the blind and sighted would suggest that the effects of blindness operate upon the developing visual pathway through the same mechanisms that shape independent and shared variation in the sighted. These questions are best addressed in large populations that maximize individual variation. We therefore collected structural magnetic resonance image (MRI) data from 53 blind and 59 normally sighted controls. The blind subjects were a heterogeneous group, ranging from those with developmental anophthalmia [9, 22] to those with a post-natal onset of blindness in adulthood. Using fully automated methods, we extracted 8 anatomical measures along the visual pathway. We examined how anatomical elements of the visual pathway vary in concert and independently in the sighted, and then tested if the same or a different pattern of variation is found in the blind.

Blindness also produces brain changes not reflected in macroscopic structure. There is a decrease in the coherence of white matter fiber tracks along the visual pathway and in the splenium of the corpus callosum [9, 23, 24] although see [22]. Changes in function are also observed, most notably the development of “cross-modal” responses, in which occipital cortical areas respond to non-visual stimulation [25, 26]. As a further test of the independence of forms of anatomical variation, we asked if individual differences in the structure of the visual pathway can account for variation in these other measures.

## Methods

### Subjects

MRI anatomical data were acquired for 59 sighted (24 men, 35 women, mean age 39 ± 14 SD) and 53 blind subjects (23 men, 30 women, mean age 44 ± 18 SD). Table 1 provides additional information for the blind group. Thirteen subjects with blindness from Leber Congenital Amaurosis were of three different genotypes (RPE65, CEP290, CRB1) and have been the subject of prior reports [27–29]. The 6 subjects with bilateral congenital anophthalmia were studied at Oxford and have been described previously [9, 22]. An additional 26 sighted subjects were studied at Oxford; their data were used only to account for systematic differences in scanner properties between the Penn and Oxford sites. An additional 13 sighted subjects were studied at the University of Pennsylvania on two separate occasions; their data was used only to examine the test / re-test reliability of the anatomical measures. Studies were approved by the University of Pennsylvania Institutional Review Board and by the Oxfordshire National Health Service Research Ethics Committee. All subjects provided written informed consent. For blind subjects, the consent forms were read aloud to the subject by the investigator, and the subject was assisted in placing a pen at the signature line.

### Magnetic Resonance Imaging

#### Overview

A T1-weighted, anatomical magnetization-prepared rapid gradient-echo (MPRAGE) image was acquired for every subject. A 3-Tesla Siemens Trio with an 8-channel Siemens head coil was used at the University of Pennsylvania, and a 3-Tesla Siemens Trio with a 12-channel coil at Oxford. A set of 8 anatomical measures along the central visual pathway was derived from the MPRAGE image from each of the 112 subjects in the study. These were measures of left and right V1 cortical thickness, left and right V1 surface area, left and right pericalcarine white matter volume, lateral geniculate volume, and optic chiasm volume. Automated techniques were used to define these anatomical regions and derive the corresponding measurements from each subject, with visual inspection of the results for quality assurance. A set of control regions were also defined for the auditory system.

Additional structural and functional MRI data were collected in a sub-set of blind and sighted subjects at the University of Pennsylvania. These additional measures were arterial spin label (ASL) perfusion, diffusion tensor imaging (DTI), and Blood Oxygen Level Dependent (BOLD) functional MRI during the presentation of auditory stimuli. Individual differences in these measures were compared to individual differences in the anatomical measures derived from the MPRAGE images.

Imaging was conducted at the University of Pennsylvania using two, slightly different protocols. After confirming for each of our imaging measures that there were no significant differences in the mean measure observed in control populations studied with the two approaches, data from the two protocols were combined. The imaging data from all subjects was visually inspected for imaging and movement artifacts. A replacement image set was acquired from subjects in whom artifacts were found.

#### Structural MR Imaging

A T1-weighted, 3D MPRAGE image was acquired for every subject. Images obtained at the University of Pennsylvania were acquired using either Protocol 1: 160 slices, 1 × 1 × 1 mm, repetition tine (TR) = 1.62 s, echo time (TE) = 3.87 ms, inversion time (TI) = 950 ms, field of view (FOV)= 250 mm, flip angle = 15°; or Protocol 2: 160 slices, 1 × 1 × 1 mm, TR = 1.81 s, TE = 3.51 ms, TI = 1100 ms, FOV= 250 mm, flip angle = 9°. Images were acquired at Oxford using: 176 slices, 1 × 1 × 1 mm, TR = 15 ms, TE = 6 ms.

**Table 1:**
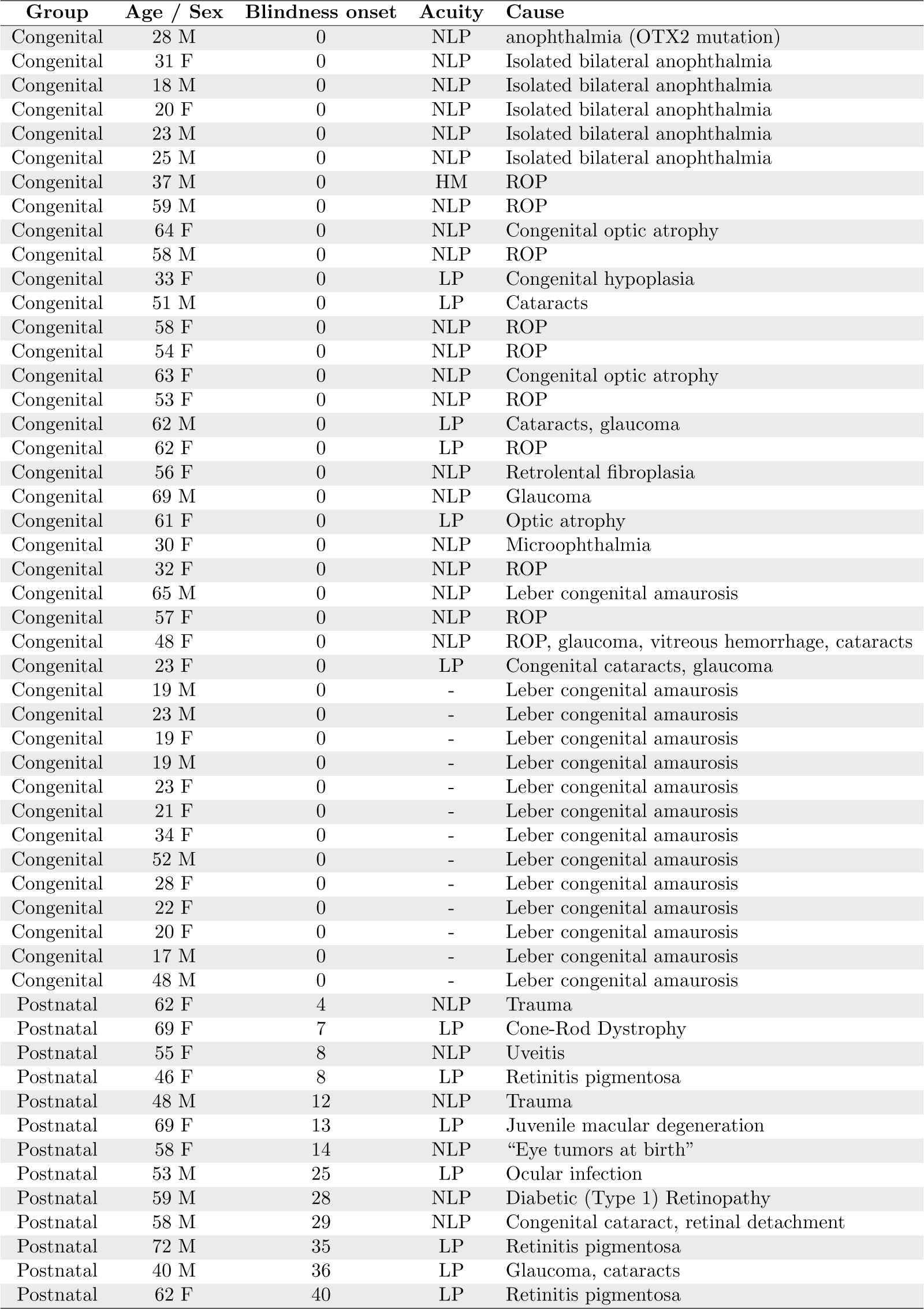
Blind subject group details. Age at blindness onset given in years. ROP - Retinopathy of Prematurity; NLP - No light perception; LP - Light perception. Values unavailable for some subjects.

#### Perfusion Imaging

An arterial spin labeled (ASL) perfusion MRI sequence was used to measure resting cerebral blood flow (CBF) in 47 subjects (16 sighted, 31 blind). These data were acquired using two different protocols. For Protocol 1, pulsed ASL was collected during a single scan of 8:08 minutes duration which consisted of 60 label/control pairs with 3.44 × 3.44 mm nominal resolution in 20 axial slices, 4 mm thickness, 1 mm gap using gradient-echo echoplanar imaging with TR = 4 sec, TE = 17 msec, TI1/TI2 = 700/1900 msec. For Protocol 2, pseudo-continuous ASL was collected during a single 5:28 minute scan which consisted of 40 label/control pairs with 2.3 × 2.3 mm nominal resolution in 20 axial slices, 5 mm thickness, 1 mm gap using gradient-echo echoplanar imaging with TR = 4 sec, TE = 60 msec, post-labeling delay time 1200 msec, and labeling time 1500 msec. Scanning was conducted while subjects rested quietly with eyes closed in darkness.

#### Diffusion Tensor Imaging

Diffusion tensor imaging (DTI) data were collected for 59 of the subjects (25 sighted, 34 blind). Each DTI scan constituted a single b=0 volume followed by either 12 (b=1000 s/mm^2^) volumes or 30 (b=1000 s/mm^2^) volumes at different gradient directions. For the DTI scans with 12 directions, the imaging parameters were: FoV = 220 mm, matrix = 128 × 128, TR = 8000 msec, TE = 82 msec, 70 slices with thickness 2 mm, interleaved. For the DTI scans with 30 directions, the imaging parameters were: FoV = 220 mm, matrix = 128 × 128, TR = 6300 msec, TE = 85 msec, 53 slices with thickness 2.1 mm, interleaved.

We note that our diffusion tensor measurements used a relatively small number of directions to allow reliable estimates of fractional anisotropy from a brief scan. These images are poorly suited for fiber tracking techniques to define and separate the geniculostriate and commisural fibers. We therefore used an anatomical region of interest approach to identify the optic radiations and the splenium of the corpus callosum and measured the fractional anisotropy within this combined region.

#### BOLD Imaging

BOLD fMRI data were collected for 52 subjects (19 sighted, 33 blind) using one of two protocols. For Protocol 1, echoplanar data were acquired in 45 axial slices with 3 mm isotropic voxels in an interleaved fashion with 64 × 64 in-plane resolution, field of view=192mm, TR=3000 ms, TE=30 ms, flip angle 90°, and 64 base resolution. Four scans, two with 100 TRs each, and two with 150 TRs each, were collected. For Protocol 2, echoplanar data were acquired in 44 axial slices with 3 mm isotropic voxels in an interleaved fashion with 64 × 64 in-plane resolution, field of view=192mm, TR=3000 ms, TR=30 ms, flip angle 90°, and 64 base resolution. Two scans with 150 TRs each were collected. Scanning was conducted with eyes closed in darkness.

#### Auditory Stimulus for the fMRI Experiment

During BOLD fMRI scanning, participants listened to spoken sentences presented in 30 second blocks, interleaved with control blocks consisting of the same sentences played in reverse or matched white noise. Each stimulus sentence had the structure “The noun is present participle” or “The noun is adjective”. A total of 150 stimulus sentences were created for the fMRI study, with 50 from each of three sensory modality categories (tactile, auditory, visual). Thirty-seven sentences in each category were considered semantically “plausible” and 13 were “implausible”. The stimuli were selected from a pool of 352 candidate stimulus sentences (88 plausible tactile, 32 implausible tactile, 88 plausible auditory, 27 implausible auditory, 88 plausible visual, and 29 implausible visual). The Kucera-Francis written frequency score of the sentences [30] were matched between modality categories. Implausible sentences paired nouns with semantically incongruent adjectives or verbs (e.g., “the chair was grunting”). Final selection of the sentences was based upon normative ratings for plausibility and semantic category assignment of each candidate stimulus. To obtain these ratings, the candidate stimuli were rated by 20 sighted control participants (who did not contribute MRI data and are otherwise not included in this study). In this process, each sentence was visually presented on a testing laptop computer. The pilot subject judged each sentence as “plausible” or “implausible”, and assigned the sentence to a sensory category: tactile, auditory, or visual. The 352 candidate sentences were presented in random sequence. We selected the stimulus sentences which received the highest consensus ratings for sensory category and plausibility. The complete list of stimulus sentences may be found online (https://cfn.upenn.edu/aguirre/wiki/public). The stimulus sentences were recorded using Microsoft Sound Recorder v 5.1 (PCM 22.06KHz, 16 bit, mono). All sentences were spoken in the same female voice. Each sentence was spoken in slightly less than 3 seconds. We used WavePad v.3.05 (NCH Swift Sound) to normalize each stimulus to the maximum auditory level and apply auto-spectral subtraction to voice. Fifty control stimuli were created by reversing the sentences in time. An additional, white noise stimulus was created. During the fMRI scans, 10 stimuli were presented in 30 second blocks with the order of blocks varied across subjects. Subjects made a button-press for each sentence that was semantically implausible, as well as for the first reverse and white-noise control stimulus in each block.

In Protocol 1, sentences from different sensory modalities were played together in a 30 second block. In Protocol 2, sentences from different sensory modalities were intermixed in blocks. Protocol 2 also presented fewer total auditory blocks.

### Image pre-processing

#### Structural MR Imaging

The FreeSurfer (v5.1) toolkit (http://surfer.nmr.mgh.harvard.edu/) [31–34] was used to process anatomical MPRAGE images from the subjects to construct white matter, pial, inflated, and spherical brain surfaces. Briefly, this processing includes spatial inhomogeneity correction, non-linear noise-reduction, skull-stripping [35], subcortical segmentation [36, 37], intensity normalization [38], surface generation [31, 32, 39], topology correction [40, 41], surface inflation [32], registration to a spherical atlas [33] and thickness calculation [34]. This approach matches morphologically homologous cortical areas based on the cortical folding patterns with minimal metric distortion and allows sampling at subvoxel resolution and detection of cortical thickness differences at the sub-millimeter level. Cortical thickness was estimated at each point across the cortical mantle by calculating the distance between the gray/white matter boundary and the cortical surface.

#### Perfusion Imaging

Image data processing and analyses were carried out with custom routines developed in Interactive Data Language (http://www.exelisvis.com/ProductsServices/IDL.aspx), which was used to quantify CBF values and reconstruct CBF maps for perfusion analyses (part of the ASLToolbox [42] (https://cfn.upenn.edu/zewang/ASLtbx.php). For each participant in Protocol 1 and 2, echoplanar images were first realigned to correct for head motion. The perfusion-weighted image series was then generated by pair-wise subtraction of the label and control images, followed by conversion to an absolute cerebral blood flow image series (ml/100g/min) based on a either a PASL perfusion model assuming a blood T1 of 1.5s at 3T for Protocol 1 data or a single-compartment CASL perfusion model for Protocol 2 data [43]. A single mean CBF image was generated for each subject and co-registered to the MPRAGE image for that subject. The CBF value at each voxel was then scaled by the global mean of the CBF image.

#### Fractional Anisotropy

Each diffusion-weighted volume was skull-striped and co-registered to the first b=0 volume using a rigid affine transformation to correct for distortion caused by eddy-current effects and simple head motion using FMRIB’s software library and diffusion toolbox v2.0 (FSL; http://www.fmrib.ox.ac.uk/fsl/). The diffusion tensor of each voxel was calculated by a linear least-squares fitting algorithm [44]. After rendering the diffusion tensor along the diagonal, the three diffusion tensor eigenvalues were obtained (mean diffusivity, axial diffusivity, and radial diffusivity). Fractional anisotropy (FA) of each voxel was derived based on the three eigenvalues. The FA was used as a measure of the degree of diffusion anisotropy. FA varies between 0, representing isotropic diffusion, and 1, in the case of the diffusion restricted to a single direction. The individual diffusion-weighted images from each subject were co-registered to subject specific anatomy in FreeSurfer using FSL-FLIRT with 6 degrees-of-freedom under a FreeSurfer wrapper (bbregister). The resulting registration matrices were used to transform the FA maps to FreeSurfer anatomical space and also to transform regions of interest (ROI) estimated in FreeSurfer space to subject-specific diffusion space.

#### BOLD Imaging

Data were sinc interpolated in time to correct for slice acquisition sequence, motion-corrected with a six parameter, least-squares, rigid body realignment using the first functional image as a reference and co-registered to the anatomical image. The fMRI data were smoothed in space with a 0.3 voxel full-width at half-maximum isotropic Gaussian kernel. Covariates of interest were modeled as a simple boxcar and convolved with a standard hemodynamic response function [45]. Covariates of no interest included global signal and “spikes”. The latter were identified by automated analysis (time points with excursions of greater than 3 standard deviations from the mean) and visual inspection and then modeled as impulses. The beta effect for each covariate (expressed as percentage signal change) was derived at each voxel. The echoplanar data in subject space were co-registered to subject specific anatomy in FreeSurfer using FSL-FLIRT with 6 degrees-of-freedom under a FreeSurfer wrapper (*bbregister*). This co-registration step allows volumetric data from each subject to be mapped to the subject’s left and right hemispheric surfaces. The volumetric beta maps of percentage signal response to “Sentences vs White Noise” and “Sentences vs. Reverse” were derived from each subject and then projected to the subject’s cortical surface.

### Regions of interest

A set of cortical and subcortical regions along the visual pathway were defined using automated methods, and subjected to visual inspection and manual correction of any errors in segmentation.

The boundary of primary visual cortex was defined for each subject by anatomical features of the cortical surface using FreeSurfer [46, 47]. The V1 region of interest in the native surface space for the subject was mapped onto the volumetric space for that subject and a binary volumetric mask was created. The pericalcarine white matter region consists of the white matter voxels that underlay primary visual cortex. This region was demarcated automatically in the native anatomical space for each subject using the Destrieux 2009 atlas segmentations [48]. The optic chiasm was identified through automated subcortical segmentation using FreeSurfer [37].

The splenium and forceps of the posterior part of corpus callosum were defined as a conjoint region. The FreeSurfer segmentation routine divides the corpus callosum into 5 equally spaced ROIs along the anterior-posterior axis; the splenium was defined in each subject as the most posterior ROI. Independently, the splenium of the corpus callosum in Montreal Neurologic Institute (MNI) space was identified using the Johns Hopkins University, DTI-based white-matter atlases [49, 50]. The splenium in MNI space technically consists of the splenium and forceps. This ROI in MNI space was warped back to individual subject anatomical space in FreeSurfer using diffeomorphic warping in the Advanced Normalization Tools (ANTs) (http://stnava.github.io/ANTs/). The union of these two regions was obtained and binarized to produce a unified region. Visual inspection was conducted for each subject to remove any voxels from the segmentation that might overlap with gray matter or the ventricles. The optic radiations were identified in MNI space using the Jülich histological atlas [51–53] and warped to individual subject space in FreeSurfer using diffeomorphic warping in ANTs. Any non-white matter voxels were removed by visual inspection. A lateral geniculate nucleus (LGN) ROI was defined within MNI space using the Jülich atlas [51–53].

A parallel set of regions were defined along the auditory pathway for a control analysis. The transverse temporal gyrus region (corresponding to primary auditory cortex) was demarcated automatically in the native anatomical space for each subject using the Destrieux 2009 atlas segmentation [48]. The transverse temporal white matter region consists of the white matter voxels that underlay transverse temporal gyrus. This region was demarcated automatically in the native anatomical space for each subject using the Destrieux 2009 atlas segmentation [48]. A medial geniculate nucleus (MGN) ROI was defined within MNI space using the Jülich atlas [51–53].

### Calculation of measures

Cortical thickness for a region was estimated separately for the left and right hemisphere using FreeSurfer, as was whole brain cortical thickness. The whole brain measure was used to adjust local thickness measures to account for differences in overall cortical thickness between individuals.

Cortical surface area for a region was defined as the sum of the area (mm^2^) enclosed by all vertices comprising the region at the gray-white matter boundary. Whole brain surface area was estimated for the left and right hemisphere using FreeSurfer. The whole brain measure was used to adjust local surface area measures to account for differences in overall surface area between individuals.

Volumes for the left and right pericalcarine white matter, transverse temporal white matter, and optic chiasm were estimated using FreeSurfer. Supra-tentorial volume was also measured and included the entire sub-pial intra-cranial volume excluding the cerebellum and brain stem. This measure was used to adjust volumetric measures to account for differences in overall brain size between individuals. Intracranial volume was also obtained and reflects the volume of all tissue (gray, white, and CSF) within the skull. This measure was also used to adjust volumetric measures to account for differences in overall brain size between individuals.

The LGN and MGN volumes were estimated using Tensor Based Morphometry (TBM). TBM calculates the Jacobian determinant for each voxel of the deformation field that relates an individual brain to the common template brain in MNI space, thus providing a measure of tissue growth or shrinkage for each voxel of the brain [54]. TBM is calculated directly from the deformation field relating all the voxels of the individual brain to the target brain in standard MNI space [55]. The mean Jacobian determinant of the deformation field was extracted for the LGN and MGN ROIs. This indirect index (as opposed to direct volumetric measurement) was used as there are no clear anatomical boundaries that define the LGN and MGN within a T1 image.

Scaled mean CBF (mean CBF/global CBF) under resting conditions was extracted from the combined left and right hemisphere V1 ROI.

The percent signal change for the contrast of “sentences vs white noise” and “reverse sentences vs. white noise” was derived for each subject from primary visual cortex.

The mean FA value in the splenium and optic radiations was obtained from the combination of these regions of interest in each subject. Derivation of the FA value from this combined region of interest obviated the need to segment the tracks attributable to the forceps of the splenium and the optic radiations.

### Reconciliation of Oxford and University of Pennsylvania data

MPRAGE images were acquired from 26 sighted controls at the Oxford site (16 men, 10 women, mean age 27 ±5 SD); data from these sighted controls at Oxford were used only for this adjustment between imaging sites and were not otherwise included in the study. Age and gender matched sighted controls at the University of Pennsylvania were identified within the entire set of 59 sighted control subjects (16 men, 10 women, mean age 27 ±5 SD). The set of 8 anatomical measures were derived from the MPRAGE images from both cohorts. The mean of each measure in the Oxford controls was then compared to the mean of each measure in the matched University of Pennsylvania sighted controls. The measures from the anophthalmic subjects studied at Oxford were then adjusted by this mean difference. This correction accounted for any small, consistent difference in the measures that might be attributed to the difference in pulse sequence or head coil between the sites. The absolute size of these adjustments was on the order of 1-2% of the mean measurement, and in all cases less than 6%. The size of these adjustments were between 3 and 30 times smaller than the mean difference in measurement size between the blind and sighted populations we observe.

### Reconciliation of Protocol 1 and 2 data from the University of Pennsylvania

Data were collected at the University of Pennsylvania under two slightly different protocols. Using unpaired t-tests, we examined if protocol differences were associated with differences in the mean of measures obtained in the sighted control group.

There was no significant difference in the eight anatomical measures derived from the MPRAGE images in the control group between the two protocols [t(57 df), all p values >0.18].

This comparison was conducted as well for the measures derived from perfusion, DTI, and BOLD fMRI. The V1 CBF measure (after scaling by global CBF) did not differ between the control subjects studied under Protocol 1 and Protocol 2 [t(14 df) = 1.69, p = 0.11].

No difference in the control subjects was found between the two DTI protocols for the average FA of splenium and optic radiations [t(23 df) = −0.69, p = 0.49].

The average BOLD fMRI response in V1 did not differ in control subjects between the data collected under Protocol 1 and Protocol 2 [t(17 df) = 0.31, p = 0.76].

Given the absence of measurable differences, we combined the data from the two protocols in subsequent analyses.

### Adjustment of anatomical measures for age, gender, and overall size effects

Each of the 8 anatomical measures were adjusted for the effects of age, gender, and variations in overall brain size within a general linear model. First, the data from all subjects (blind and sighted) were combined. Covariates were created that reflected gender, age, and a quadratic and cubic polynomial expansion of age. A covariate that modeled the effect of data being collected from Protocol 1 or Protocol 2 was also included to ensure that no main effect of this factor could influence the subsequent analyses.

Covariates were also included to capture overall scaling effects of brain size. For measures of cortical thickness, this covariate was the mean global cerebral cortical thickness. For measures of surface area, this covariate was total cerebral surface area. For measures of volume, a measure of total supratentorial volume and a measure of total intracranial volume was included. Each of these size measures were also expanded to reflect quadratic and cubic effects of the variable. After fitting, the residual data from this model was taken as the adjusted measure. For every measure, significant variance was accounted for by the linear effect of a size covariate; there was also a quadratic effect of intracranial volume on the size of the optic chiasm. Gender and age (and the polynomial expansion of age) did not independently explain significant variance in the measures, with the exception of a modest quadratic effect of age upon left hemisphere V1 surface area, and a cubic effect of age upon left hemisphere pericalcarine volume.

We next tested if there are further effects in the data that manifest as an interaction of group (blind or sighted) with adjustment factor (age, gender, overall size). We modeled these interaction terms (and their cubic and quadratic polynomial expansions) and found that they did not explain significant variance in the data for any of the 8 measures (all p-values >0.15). Therefore, the data were not adjusted for these interaction effects.

### Hierarchical clustering analysis

We conducted a data reduction step to combine the 8 anatomical measures derived from the MPRAGE images into fewer dimensions. This analysis was conducted separately for the data from each group (sighted and blind). First, the measures were z-standardized within group. To do so, the mean value of a measure across the group (sighted or blind) was subtracted from the measurement for each subject, and then the set of values for a given measurement was divided by the standard deviation of a given measurement across the group. The motivation for this transformation of the data is to allow each measurement to be equally weighted in the subsequent clustering analysis. The resulting data matrix (59 × 8, subjects × measures for the sighted group; 53 × 8 for the blind) was submitted to the MATLAB hierarchical clustering analysis routines of *pdist* and *linkage*, using a Euclidean distance metric and Ward’s minimum variance criterion for clustering. The ability of the resulting dendrogram to model the pair-wise distances in the data was evaluated by calculating the cophenetic correlation coefficient using the MATLAB function *cophenet*.

The reliability of the assignment of anatomical measures to clusters was examined with a bootstrap re-sampling analysis. The set of blind (or sighted) subjects was sampled with replacement and the cluster analysis repeated. We measured the proportion of the 1000 re-samples for which the same anatomical measures were assigned to a given cluster.

**Table 2:** Mean group differences in the anatomical measures. The anatomical measurements for each group are given, after adjustment for individual differences in brain size and removal of age and gender effects. The Cohen’s *d* is the difference in means relative to the standard deviations. LH - Left Hemisphere; RH - Right Hemisphere; V1 - Primary Visual Cortex; LGN - Lateral Geniculate Nucleus.

| | Measure [units] | mean ±SD Sighted | mean ±SD Blind | Cohen's $d$ |
| --- | --- | --- | --- | --- |
| 1 | LH V1 cortical thickness [mm] | 1.64±0.09 | 1.76±0.20 | -0.8 |
| 2 | RH V1 cortical thickness [mm] | 1.69±0.10 | 1.75±0.16 | -0.4 |
| 3 | LH V1 surface area [mm<sup>2</sup>] | 2529±298 | 2183±325 | 1.1 |
| 4 | RH V1 surface area [mm<sup>2</sup>] | 2382±355 | 2007±292 | 1.2 |
| 5 | LH Pericalcarine volume [mm<sup>3</sup>] | 3302±578 | 2623±592 | 1.1 |
| 6 | RH Pericalcarine volume [mm<sup>3</sup>] | 3332±562 | 2576±623 | 1.2 |
| 7 | Optic chiasm volume [mm<sup>3</sup>] | 250±69 | 178±60 | 1.1 |
| 8 | LGN volume [log Jacobian] | 0.94±0.12 | 0.78±0.13 | 1.3 |
| | Whole brain cortical thickness [mm] | 2.53±0.13 | 2.53±0.13 | 0.00 |
| | Cerebral surface area [meters<sup>3</sup>] | 0.18±0.02 | 0.17±0.02 | 0.08 |
| | Supratentorial volume [liters] | 1.02±0.11 | 1.00±0.13 | 0.04 |
| | Intracranial volume [liters] | 1.35±0.28 | 1.25±0.27 | 0.09 |

## Results

### Mean differences in visual pathway anatomy between the blind and the sighted

Measures of visual pathway anatomy (Fig 1A) were derived from an MPRAGE image of the brain for 59 sighted and 53 blind subjects. The measures were adjusted to remove the effect of individual variation in overall brain size and surface area, and the effects of variation in age and gender. Separate measures of left and right cortical structures were obtained as the effects of blindness can be lateralized [56].

As would be expected from many prior studies, there were highly significant and substantial differences between the blind and sighted groups in the mean of each measure (Table 2). The relative area and volume of anatomical structures along the visual pathway are greater in the sighted, with the exception of the thickness of the gray matter layer of V1 cortex which is increased in the blind. In unpaired t-tests (110 df) between the groups, these differences were all significant at the p=0.00025 level or lower, with the exception of the difference in V1 cortical thickness in the right hemisphere, which had an associated p-value of 0.027. Table 2 also presents the Cohen’s *d* score for these group comparisons, indicating the size of the group difference relative to the standard deviation across individuals.

The effect of individual variation in overall brain size was removed from the measures. We nonetheless checked if substantial differences in these whole-brain measures exist between the two groups. The lower portion of Table 2 provides the group means for each of the whole-brain indices used to adjust the local visual pathway measures. The differences between groups were small relative to individual variation, as indicated by the Cohen’s *d* scores which were all less than 0.1.

We considered the possibility that the effects we find along the visual pathway reflect general mechanisms of brain development that are altered in a non-specific way in our blind populations. We repeated our analyses using seven analogous anatomical measures from the auditory pathway. We found minimal difference between the blind and sighted groups for the seven measures (Cohen’s *d* for all measures between 0 and 0.4), supporting the anatomical specificity of our primary findings.

**Figure 1:**
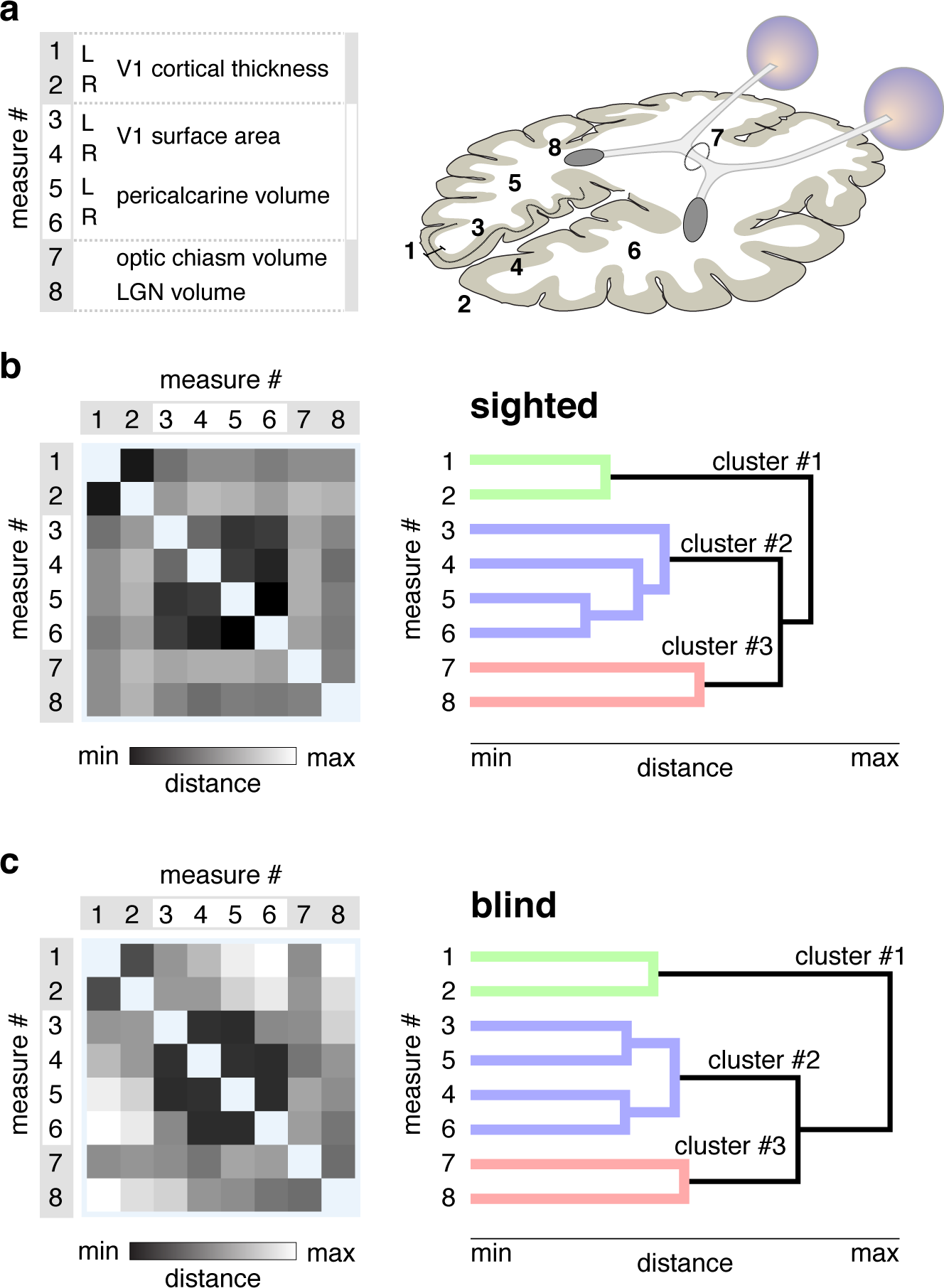
Patterns of shared variation in visual pathway anatomy. *(A)* The eight measures of visual pathway anatomy are illustrated on an axial schematic of the human brain. The groupings of the measures are to assist subsequent interpretation of the data. *(B)* The Euclidean distance matrix and dendrogram for the 8 measures across the sighted population. *Left*. The square-root, sum-squared difference in values between two measures across subjects provides a measure of Euclidean distance. Darker shades indicate pairings of measures that have similar variation across subjects, and thus lower distance values. *Right*. The distance matrix was subjected to hierarchical clustering, yielding a dendrogram. The length of each branch reflects the distance between the paired measures. The three primary clusters of anatomical variation are colored green, blue, and red. *(C) Left*. The distance matrix across the 8 measures for the blind population. A similar overall structure is seen as compared to the sighted. *Right*. The dendrogram derived from measures from the blind subjects. The same overall cluster structure is seen. Note that there is some rearrangement in the measurements assigned to cluster #2 in the blind as compared to the sighted.

### Correlated anatomical variation in visual pathway anatomy

While there is significant separation in the group mean of measures of visual pathway anatomy, there is also broad variation about the means within each group. This individual variation within a group (blind or sighted) is roughly equal to the size of the difference between groups, as indicated by Cohen’s *d* scores that are close to one.

This variation cannot be attributed to the effects of gender, age, or individual differences in overall brain size or cortical thickness, as these effects were modeled and removed from the data. We confirmed that the variation is not the simply the result of measurement noise. We collected an MPRAGE brain image from an additional thirteen, normally sighted subjects (not otherwise included in the study) on two occasions between 7 and 194 days apart. We derived the eight measures of visual pathway anatomy and compared the measures obtained on the two occasions. Across the subjects, individual variation in the measures was strongly correlated across the two imaging sessions (Pearson correlation values between the first and second measurement session *>*0.77 for all measures, and >0.90 for six of the eight measures).

As this variation seems likely to reflect true individual differences, we next asked if there are consistent patterns of variation in the relative size of visual pathway components across individuals. For example, in previous studies of normally sighted people, occipital lobe surface area and volume were found to be correlated, but relatively independent of V1 cortical thickness [19]. We examined this first in data from the sighted subjects. The set of 8 measurements from 59 sighted people was submitted to a hierarchical clustering analysis. The resulting Euclidean distance matrix and dendrogram shows the structure of correlation across subjects in variation in the anatomical measures, and the clustering of measures into covarying groups (Fig 1B). This dendrogram had high explanatory power in the data, with a cophenetic correlation coefficient of 0.88 (the maximum possible being 1.0).

The dendrogram identifies three primary clusters of anatomical variation across subjects. Similar to prior findings [19], cortical thickness in left and right V1 are highly correlated. This first cluster is relatively independent from a second cluster, composed of the surface area and volume of the left and right V1 cortex and pericalcarine white matter. The size of the optic chiasm and lateral geniculate nucleus form a third and final cluster. These clusters of measures were a stable property of the data. We conducted a bootstrap analysis of the data, and found the same anatomical measures were assigned to the first, second, and third clusters in 99, 98, and 79% of the re-samples, respectively.

We next asked if the same pattern of visual pathway variation is found in the population of blind subjects. It is possible that blindness would cause some anatomical structures that are normally independent to become correlated, perhaps through a shared mechanism of atrophic change. Instead, a notably similar clustering of anatomical variation was found in the blind as was seen in the sighted (Fig 1C). This structure was a stable property of the data (cophenetic correlation = 0.82; replicability of clusters 1, 2, and 3 was 100, 99, and 70%, respectively).

These findings indicate that there are consistent patterns of individual variation in brain structures along the visual pathway, and that the form of this variation is highly similar in the sighted and blind. The clustering of anatomical variation also indicates that the set of 8 measures can be combined into three composite measures. As the units differ substantially between the measures, we z-transformed each measure after combining the blind and sighted populations, so that the mean value across all subjects is zero, with positive values being typical of sighted subjects and negative values typical of blind subjects. In this representation of the data, the measurement of V1 cortical*thickness* is reversed to become a measurement of relative*thinness*, with a thinner cortex being more typical of the sighted population. We then averaged the z-transformed measures in each of the three clusters, allowing us to summarize visual pathway anatomy for each subject with a set of three values, one for each cluster (Fig 2).

As would be expected, the mean value for each cluster differed significantly between the blind and the sighted population (unpaired t-test, cluster 1: p=0.0013; cluster 2: p <0.00001; cluster 3: p <0.00001). We then considered how heterogeneity in our population of blind subjects might be related to these anatomical measures. At a coarse level, the blind population (Table 1) can be divided into those with vision loss at birth (congenital), and those who developed vision loss after birth (postnatal). It is possible, for example, that anatomical variation could be established in utero, with no role for post-natal input upon relative size. If this were the case, the relative size of a cluster of anatomical structures would not differ between the sighted and postnatally blind. We examined the anatomical score within each cluster for the sighted, congenitally blind, and postnatally blind. For all three clusters of visual pathway anatomy, measurements from the postnatally blind were intermediate to those from the congenitally blind and sighted, although these differences were significant only for the second and third clusters.

**Figure 2:**
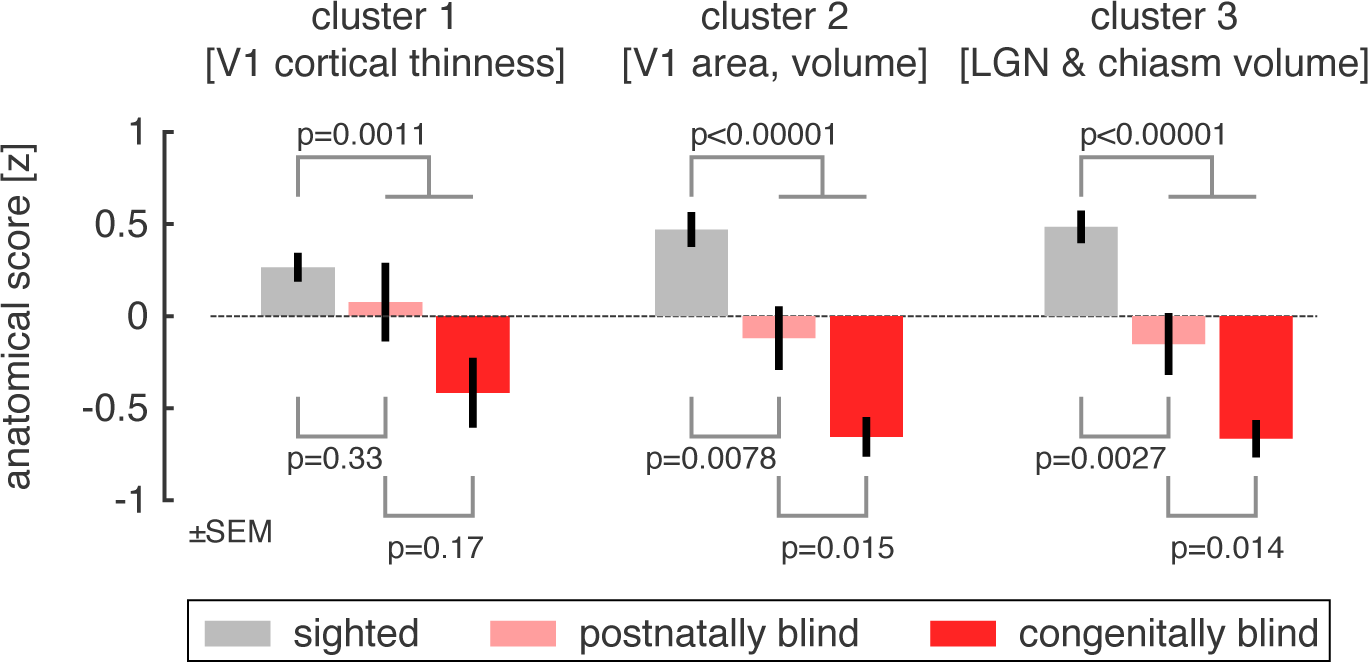
Group differences in anatomy, organized by cluster. Each anatomical size measurement was transformed to a mean-centered z-score and then averaged within a cluster. The sign of values in cluster #1 was reversed, so that positive values represent*thinner* V1 cortex. Anatomical scores within each cluster were then averaged within subject groups, corresponding to the normally sighted, postnatally blind, and congenitally blind. Across all three clusters, a graded change in the anatomical scores is seen for these subject groupings, although the difference between postnatally and congenitally blind is not significant for cortical thinness (cluster #1).

Given these findings, one might consider the possibility that the degree of visual pathway alteration is related to the age at which vision was lost, as has been seen for cortical thickness in a large population of blind subjects [10]. While an appealing idea in principle, it is difficult to test in practice as many causes of blindness in our cohort are progressive, preventing the identification of a clear age of onset. Nonetheless, we identified 13 subjects with a discreet timing of postnatal onset of blindness. We did not find a significant Spearman’s rank correlation between the reported age of blindness onset and either the eight anatomical measures, or their grouping into three clusters (all p-values >0.17).

### Group differences in white matter coherence and cross-modal responses

Given our finding of independent patterns of anatomical variation in the sighted and the blind, we next explored if these clusters of variation are differentially related to other structural and functional imaging measures. About half of our blind and sighted subjects contributed additional MRI measurements. We first tested for group (blind vs. sighted) differences in measurements of cerebral blood flow, cross-modal response, and white matter fractional anisotropy.

We obtained a measure of resting-state (eyes closed, darkness) cerebral blood flow within primary visual cortex using arterial spin-labeled MRI. Prior studies have found an increase in resting functional activity in the visual cortex of the blind as compared to the sighted (as reflected in resting glucose utilization [57]). We did not find a difference in V1 CBF between the blind and sighted groups [mean relative V1 CBF in the sighted: 1.20, and blind: 1.15; unpaired t-test (45 df) = 0.50, p = 0.62]. We tested for a difference between the congenitally and postnatally blind, as a prior study of glucose utilization had found opposite effects upon resting metabolic activity depending upon the timing of blindness [58], but did not find a significant difference [mean relative V1 CBF in the congenitally blind: 1.02, and postnatally blind: 1.22; unpaired t-test (29 df) = −1.48, p = 0.15]. As there was no group difference in this measure, we do not further consider it here.

Our subjects also underwent functional MRI scanning. Blood oxygen level dependent (BOLD) fMRI was used to measure “cross-modal” neural responses evoked within striate cortex during an auditory semantic judgment task. Measurements from primary visual cortex were obtained while subjects listened to sentences played forwards and sentences played in reverse, as compared to listening to white noise. In agreement with multiple prior studies, we found a greater response to these stimuli in the blind subjects as compared to the sighted [mean % BOLD signal change in the sighted: 0.89, and blind: 2.25; unpaired t-test (50 df) = −2.03, p = 0.048]. There was no significant difference in response between the congenitally blind as compared to the postnatally blind [t(31 df) = −1.40, p = 0.17]. We also tested for group effects in the differential V1 response evoked by forward as compared to reverse sentences, reflecting semantic content while controlling for low level auditory features. There was no significant difference between the groups in this measure [mean % BOLD signal change difference between forward and reverse sentences in the sighted: 0.59, and blind: 1.22; unpaired t-test (50 df) = −1.22, p = 0.26]. In analyses described below, we examine the relationship between cross-modal response and individual differences in anatomy. As the group difference for cross-modal response was largest for all auditory stimuli (forwards and reverse speech combined), we focused upon this measure in subsequent tests.

Finally, we obtained diffusion tensor imaging (DTI) measures of the fractional anisotropy (FA) of the optic radiations and splenium of the corpus callosum. FA values were significantly reduced in these white matter components of the central visual pathway in the blind as compared to the sighted [mean FA value in the sighted: 0.48, and blind: 0.44; unpaired t-test (57 df) = 3.29, p = 0.0017]. This measure was not different between the congenitally and postnatally blind groups [t(32 df) = 1.10, p = 0.28]. We may have lacked the statistical power needed to replicate prior findings of a difference in these subgroups [59].

### Anatomical size variation is related to white matter coherence and cross-modal responses

We observed earlier that individual differences in the size of structures along the visual pathway can be grouped into three primary clusters of variation, and that this clustering is similar in the sighted and blind groups. We next asked if anatomical size variation is related to individual differences in cross-modal responses and white matter organization in the blind and sighted. The average score for each of the three anatomical clusters was obtained for each subject. Within a linear model, we then tested if the anatomical scores could be used to predict either cross-modal responses or fractional anisotropy measures.

Anatomical variation was significantly related to the degree of cross-modal response within area V1 across subjects [F(3,54 df) = 9.54, p = 0.00028] (Fig 3A). We examined the weights on the model (Fig 3B), and found that the relationship between visual pathway anatomy and cross-modal BOLD fMRI response is mediated primarily by cortical thickness. Across subjects, the thicker V1 cortex, the greater the cross-modal response. The two other clusters of anatomical variation were not significantly related to cross-modal activity.

Anatomical variation in the visual pathway was also significantly related to the fractional anisotropy measured from the optic radiations and splenium [F(3,56 df) = 4.45, p = 0.016] (Fig 3C). In this case, it was variation in the third anatomical cluster—reflecting the relative size of the optic chiasm and lateral geniculate nucleus—that was related to fractional anisotropy of the visual pathway (Fig 3D). The larger the optic chiasm and lateral geniculate nucleus, the higher the FA score measured in the optic radiations and splenium.

**Figure 3:**
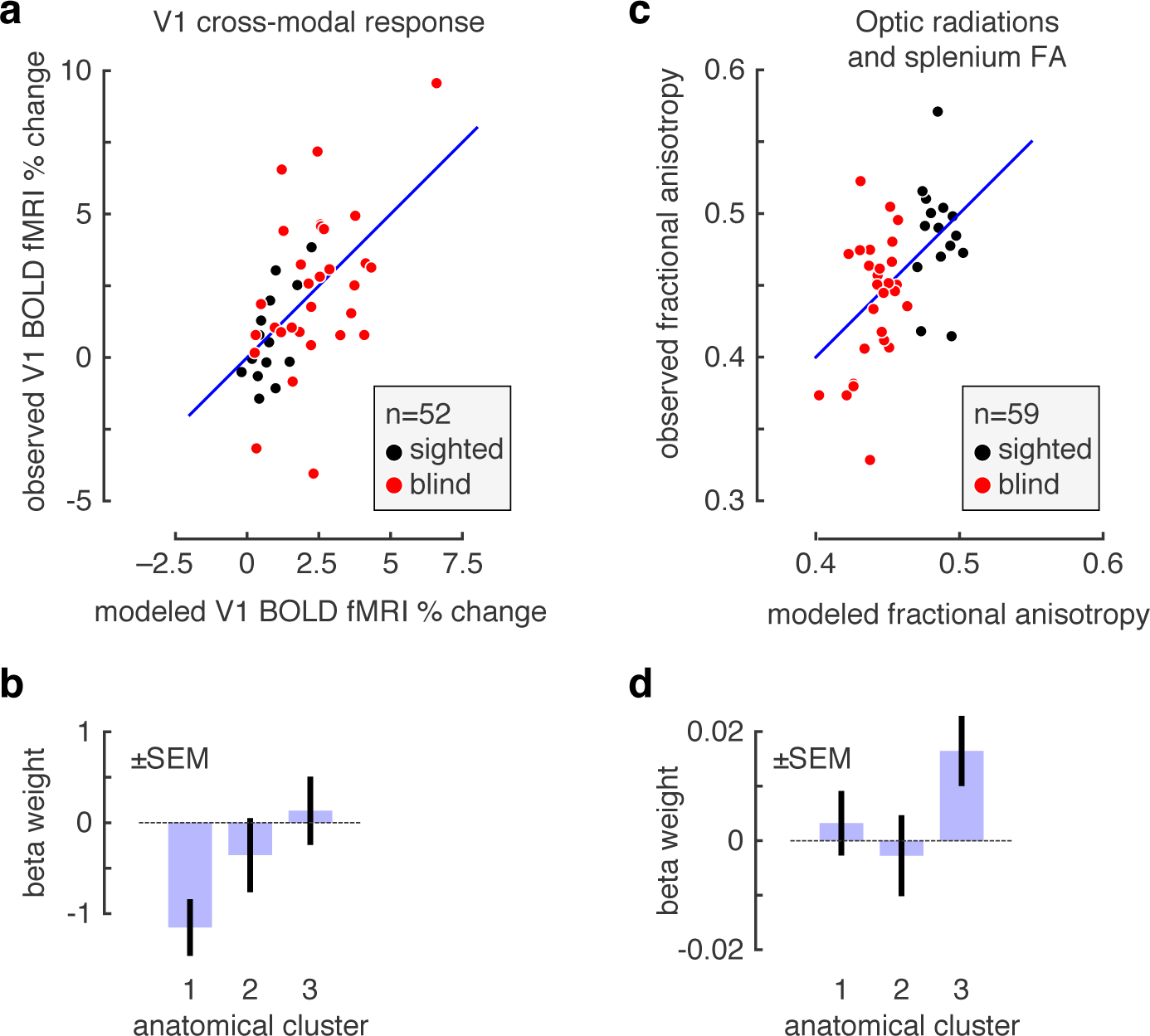
Relation of clustered anatomical variation to cross-modal response and fractional anisotropy. *(A)* For each of 52 subjects (blind and sighted), we obtained the BOLD fMRI response in V1 while subjects listened to auditory sentences played forwards and in reverse, as compared to white noise. We modeled the ability of individual variation in the three anatomical clusters to account for variation in cross-modal BOLD fMRI response. For each subject, the x-axis gives the prediction of the model for BOLD fMRI response, and the y-axis the observed response. There was a significant model fit (p=0.00028). *(B)* Model weights for the fit to the cross-modal response data. Shown are the mean and standard error of weights upon each of the clusters of anatomical variation in their prediction of V1 BOLD fMRI response. Only the first cluster of anatomical variation (V1 cortical*thinness*) had a fitting weight significantly different from zero. The loading on this weight is negative, indicating that*thicker* V1 cortex predicts greater cross-modal responses. *(C)* For each of 59 subjects, we measured fractional anisotropy within the optic radiations and splenium of the corpus callosum. We modeled the ability of individual variation in the three anatomical clusters to account for variation in FA. For each subject, the x-axis gives the prediction of the model for FA, and the y-axis the observed measure. The entire model fits the data above chance (p=0.016). *(D)* Model weights for the fit to the FA data. Shown are the mean and standard error of weights upon each of the clusters of anatomical variation in their prediction of the FA measure. Only the third cluster of anatomical variation (chiasm and LGN volume) had a fitting weight significantly different from zero.

These correlations are driven in part by group effects. That is, both the anatomical size measures, and the measures of cross-modal activity and fractional anisotropy, contain a mean difference between the blind and sighted subjects. We examined if anatomical variation along the visual pathway could predict cross-modal activity and fractional anisotropy independently of group effects. After modeling and removing the group (blind vs. sighted) effect, we find that the first anatomical cluster continues to predict cross-modal response (p=0.0015). In a post-hoc test, we confirmed that a positive Pearson correlation is present between cortical thickness and cross-modal BOLD fMRI response for both the sighted (r=0.41) and the blind (r=0.45). In contrast, after accounting for the group effect, there was no longer significant relationship between the third anatomical cluster and the DTI measures (p=0.087).

In sum, individual differences in macroscopic anatomical structure in the blind and sighted are correlated with individual differences in other measures of brain structure and function. Importantly, these relationships are separable. Area V1 cortical thickness is related primarily to cross-modal responses at this site, while the size of the optic chiasm and lateral geniculate are related to fractional anisotropy within the optic radiations and splenium. In the case of cross-modal responses, individual variation in cortical thickness is predictive of BOLD response independently of the group effect of blindness.

## Discussion

Our study emphasizes the graded nature of structural brain changes that accompany human blindness, and places them within the context of normal variation in the sighted. Individuals vary substantially and meaningfully in the relative size of the anatomical components of the central visual pathway. In normally sighted people, we find the thickness of V1 gray matter is largely independent of the surface area and white matter volume of V1 cortex, and in turn independent of the relative size of the optic chiasm and lateral geniculate nucleus. Large, cross-sectional studies of normally sighted people have previously reported that occipital lobe surface area and volume are relatively independent from cortical thickness [19, 20, 60], and these anatomical features have opposite relationships with neural and perceptual measures of visual discrimination [61].

We find these same three clusters of variation in the visual pathway of blind people. It need not have been so. Blindness could have reconfigured the variation of visual pathway anatomy, causing, for example, linked atrophy across structures that are normally independent. Instead, the effect of blindness is to extend the range of variation seen in normal subjects along each of the three dimensions of visual pathway anatomy. Interestingly, these effects are seen in the postnatally blind, as well as in the congenitally blind.

### Independent anatomical effects of blindness

Across blind participants, there was a varying degree of reduction in the surface area of V1 and the corresponding white matter volume of visual cortex. In the sighted, the spatial extent of area V1 is correlated with visual acuity [17] and perceptual discrimination [61]. A recent study of patients with juvenile and adult onset macular degeneration found roughly equal reductions in the volume of the optic radiations in both groups [62], consistent with our finding that this anatomical change can arise in adulthood.

Separately, there is linked variation in the size of the optic chiasm and lateral geniculate nucleus. Variation on this anatomical dimension was correlated with a DTI measure of fractional anisotropy within the optic radiations and splenium. In cross-sectional studies of children, age related increases in the fractional anisotropy of regional white matter were found to be correlated with increases in the associated volume of the region [63]. A similar mechanism may link visual pathway size and the coherence of white matter.

Finally, we find that V1 cortical thickness varies independently from these other anatomical dimensions. Early synaptic remodeling in response to visual input shapes visual cortex [11], and has been proposed as the mechanism that produces cortical thinning with development [63]. We find further that cortical thickness is related to the amplitude of cross-modal response in striate cortex. A notable feature of this finding is that it was present in both sighted and blind subjects. A relationship between cortical thickness and cross-modal responses may reflect relative preservation of early, exuberant synaptic connections that are pruned as the cortex thins during normal visual development [64]. Complementary findings are that individual differences in occipital gray matter across blind subjects are correlated with the duration of blindness, and in turn correlated with behavioral performance on non-visual tasks [56, 65], and that cross-modal responses in post-natal blindness are proportional to the loss of the visual field [66]. Our result is in disagreement, however, with a prior study of twelve people with blindness before the age of two [67]. In this prior work, a negative correlation was found between BOLD fMRI response during a sound identification task and cortical thickness in the left collateral sulcus (and at other sites). We do not have an explanation for this difference in results, but consider it possible that methodological differences in correction for age and head size effects may contribute.

Changes in visual cortex thickness have been described in postnatal causes of vision loss that involve specific portions of the visual field, although there are conflicting reports as to the direction of the effect. Young adult carriers of a mitochondrial DNA mutation causing Leber hereditary optic neuropathy have a loss of macular retinal ganglion cells and thicker visual cortex gray matter [68]. In contrast, older people with progressive retinal damage from glaucoma or macular degeneration have thinning in the corresponding eccentric locations of visual cortex [69–72]. It seems the response of the cortex to retinal damage is influenced by the spatial distribution of this loss, the age at which it occurs, or both. Earlier studies that reported gray matter loss in blinding diseases are difficult to relate to our study as a voxel based morphometry approach was used which combines changes in thickness and surface area [73].

### Limitations and alternatives

While a gradation of anatomical alteration between the congenitally blind, postnatally blind, and sighted is a feature of our results, we note that there is evidence in support of categorical, qualitative differences between the blind and sighted, and between the early and later blind [74]. For example, temporary inactivation of the occipital cortex in the blind has qualitatively different effects depending upon the age of blindness onset [21, 75].

We make use of standard techniques to segment the gray and white matter tissue compartments, and to subsequently measure cortical thickness [37]. This measurement can be influenced by the degree of intra-cortical myelination [76] and it is therefore possible that a reduction in intra-cortical myelination is the cause of what would be an apparent and not actual increase in occipital cortex thickness in the blind. We note, however, that in our data the measure of cortical thickness is independent of both the relative volume of the optic radiations and the fractional anisotropy of this structure, inconsistent with the alternative account. Further, work by others [65] has shown that the effect of blindness upon myelin content is largely confined to the peri-cortical white matter. Regardless, a difference in mechanistic interpretation does not alter the general observations that we make regarding the relationship of the measure of cortical thickness to age of blindness and cross-modal plasticity.

We examined the size and properties of the visual system between the optic chiasm and the primary visual cortex. The cortical visual system extends far beyond this point, and blindness has been found to alter both the properties of these higher visual areas and the connections of visual cortex to other brain regions [25, 77, 78]. We regard the relationship between the anatomical variation we have measured and the properties of higher visual areas and their connections as a promising area for future study. As an example of this direction of investigation, recent work finds that the covariance of early visual cortex thickness with dorsal visual areas is related to enhanced sensory abilities in the blind [79], and that there are differential effects of blindness upon dorsal and ventral stream white matter anisotropy [80]. Decoupling of occipital and frontal anatomical features has been observed in the blind [79], suggesting that this is a promising avenue of investigation. Similarly, our study examined changes along the retinofugal visual pathway, and not the brainstem system for vision that includes the superior colliculi. Recent work has shown that, while macroscopic superior colliculus structure is not altered in the congenitally blind [1], there are changes in its functional properties [81].

Our group of blind subjects varied on many clinical features. Indeed, a key feature of our study was the inclusion of a heterogeneous population of blind subjects, allowing us to leverage this variance to examine anatomical patterns. We examined the effect of the timing of the onset of vision loss (congenital or postnatal), and observed a graded degree of anatomical alteration. While other divisions of the postnatal group by age could be considered, there is evidence that these effects are better considered as a continuous variable [10]. Other aspects of blindness could also contribute to anatomical variation. Differences in braille reading, light sensitivity, visual fields, or involvement of the retinal ganglion cells in the ophthalmologic injury could influence post-chiasmal visual anatomy in ways the current study was not designed to detect. We note as well that the heterogeneity of our post-natal group could reduce our ability to detect mean differences in the blind and sighted in some measures, such as V1 CBF.

## Conclusion

To a first approximation, differences between the blind and the sighted have been found for every structural and functional measure obtained along the pathway from optic nerve to visual cortex. Our study, along with other recent work (e.g., [67, 79]), demonstrates that these numerous brain changes can be assembled into meaningful sets, and reflect extensions of the variation seen in normal development. An exciting avenue for further investigation is if particular clinical properties of blindness produce more or less alteration of the different anatomical clusters of variation that we find here.

